# Structures of TASK-1 and TASK-3 K2P channels provide insight into their gating and dysfunction in disease

**DOI:** 10.1101/2024.08.05.606641

**Authors:** Peter-Rory Hall, Thibault Jouen-Tachoire, Marcus Schewe, Peter Proks, Thomas Baukrowitz, Elisabeth P. Carpenter, Simon Newstead, Karin E.J. Rödström, Stephen J Tucker

## Abstract

TASK-1 and TASK-3 are pH-sensitive Two-Pore Domain (K2P/*KCNK*) K^+^ channels. Their functional roles make them promising targets for the treatment of multiple disorders including sleep apnea, pain and atrial fibrillation. Rare genetic mutations in these channels are also associated with neurodevelopmental and hypertensive disorders. A recent crystal structure of TASK-1 revealed a lower ‘X-gate’ that is a hotspot for missense gain-of-function mutations associated with DDSA (Developmental Delay with Sleep Apnea). However, the structural basis for gating in TASK channels and how they sense extracellular pH to regulate gating have not been fully elucidated. Here, we resolve structures for both the human TASK-1 and TASK-3 channels by cryoEM, as well as for a recurrent TASK-3 variant (G236R) associated with *KCNK9* Imprinting Syndrome (formerly referred to as Birk-Barel Syndrome). Combined with functional studies of the X-gating mechanism, these structures not only provide evidence for how a highly-conserved gating mechanism becomes defective in disease, but also provide further insight into the pathway of conformational changes that underlie the pH-dependent inhibition of TASK channel activity.

## Introduction

Two-Pore Domain K^+^ (K2P) channels are a structurally distinct subset of K^+^ channels where each gene encodes a subunit with two pore-forming domains that co-assemble as a dimer to create a single pseudotetrameric K^+^-selective pore which spans the cell membrane^1^. Originally described as ‘leak’ channels, K2P channels are now known to be regulated by many diverse stimuli, including multiple G-protein coupled receptor (GPCR) pathways^2^ thus enabling them to integrate a range of neuronal, metabolic and cellular signaling pathways into changes in cellular electrical activity^3, 4^.

In humans, K2P channels are encoded by 15 separate genes (*KCNK1-KCNK18)* and whilst most assemble as homomeric channels, several are able to coassemble as heteromeric channels with unique functional properties^5, 6^. *KCNK3* encodes the TASK-1 K2P channel and is expressed in many chemosensitive regions of the central nervous system involved in the regulation of breathing, as well as in the lung, heart, carotid body, adrenal gland and pulmonary arterial smooth muscle^1, 7, 8^. Loss-of-function (LoF) mutations in TASK-1 cause an inherited form of hypertension (Pulmonary Arterial Hypertension; PAH)^9, 10^, whilst *de novo* gain-of-function (GoF) mutations have recently been shown to underlie a separate disorder termed Developmental Delay with Sleep Apnea (DDSA)^11^. The expression of TASK channels in cells and tissues involved in both the control of respiratory drive and in mechanical ventilation, as well as its genetic association with DDSA, have therefore implicated TASK-1 as a target for the treatment of sleep apnea, and in a recent randomized, blinded, clinical trial a novel TASK channel inhibitor was found to reduce sleep apnea severity in patients with severe obstructive sleep apnea^12^.

The related TASK-3 channel is encoded by *KNCK9* and shares 54% overall amino acid identity with TASK-1 and 80% identity within the pore-forming transmembrane regions. TASK-3 is also expressed widely throughout the CNS and several peripheral tissues. Furthermore, in places where both TASK-1 and TASK3 are coexpressed, e.g., in the carotid body, heteromeric TASK- 1/TASK-3 channels appear to coassemble to form the physiologically relevant channel subtype in those cells^5, 6, 13^.

*KCNK9* Imprinting Syndrome (KIS) is a rare neurodevelopmental disorder, previously referred to as Birk-Barrel syndrome. KIS is primarily characterized by LoF mutations in the maternal copy of the gene, with the majority of patients possessing a single G236R mutation^14^. More recently, GoF mutations have also been associated with this disorder^15^, but the underlying molecular mechanisms behind this genotype-phenotype correlation remain unclear and can often be complicated by imprinting and/or mosaicism. Interestingly, TASK-3 has also been shown to be expressed in certain nociceptive sensory neurons and TASK-3 activators have been shown to display potent analgesic effects in a variety of rodent pain models suggesting this channel as a potential target for the treatment of pain^16^.

In a recent study, we determined an X-ray crystal structure of TASK-1 in complex with a compound class of TASK-1 inhibitors developed for the treatment of sleep apnea^17^. This structure also revealed several unique features of TASK-1, including a lower ‘X-gate’, a structural motif involving constriction of the M4 helices that controls opening and closing of the channel pore. The X-gate and surrounding region also appears to be a hotspot for GoF mutations underlying DDSA by creating channels which have both a higher open probability and which are resistant to GPCR-mediated inhibition^11^.

The extracellular H^+^ sensitivity of TASK channels is also an important and physiologically relevant feature with their name originally derived from their acid sensitivity^8^. TASK-1 and TASK-3 channels have pKa values of ∼7.4 and 6.7 respectively thus permitting their activity to be finely tuned by changes in extracellular H^+^ concentrations within the physiological range^18^. Previous studies have identified the pH-sensor as H98 positioned adjacent to the first selectivity filter (SF) motif (TIGYGH)^19^. The first insights into the mechanism of gating have been provided by cryoEM structures of the related TWIK-1 channel which also possesses a histidine pH-sensor in a similar position within the filter motif^20^ and more recently by a structure of TASK-3^21^, but the precise mechanism by which protonation of H98 influences channel gating in TASK channels has not been fully elucidated. It is known that even relatively small structural rearrangements within the pore-forming loops control a ‘C-type’ or inactivation gate within the SF of many K2P channels and similar changes may therefore also be sufficient to regulate gating in TASK channels^22–25^.

However, for any such SF gate to function, the lower X-gate must also open, and in many K^+^ channels such gates are allosterically coupled^26–28^. In TASK-1 we have previously shown that mutations which affect X-gating can also influence activation by external H^+^, but that they do not prevent inhibition. This suggests that if such coupling exists between these gates then it may be relatively weak^11^. However, without more detailed structural or functional information, such mechanisms remain difficult to determine. Likewise, without more structural information, the effects of disease-causing mutations remain difficult to interpret as well as impacting the rational design of better therapeutics that target these channels.

To address these issues, we have used single particle cryoEM to solve the structure of TASK-3 with and without the most frequent KIS mutation (G236R) as well as a structure of TASK-1, and we use functional studies to demonstrate that the X-gating mechanism is conserved in TASK-3. Importantly, these structures reveal how the pH-sensor in TASK-1 channels begins to move in response to protonation to control channel gating, and how the X-gate may also begin to open. We discuss how these movements relate to other structures of TASK-3 recently resolved by cryoEM at different pH values^21^ and how these various new structures provide further insight into the gating mechanism of TASK channels and their associated channelopathies, as well as expanding the structural landscape of TASK K2P channels as druggable targets.

## Results and Discussion

### CryoEM Structure of TASK-1

The previous crystal structure of TASK-1 revealed a lower X-gate which was also seen in the two inhibitor-bound structures resolved in that study^17^. However, the contacts required for crystallisation occasionally influence structures to produce non-native conformations. An advantage of single particle cryoEM is that additional native conformations can often be resolved. We therefore determined a cryoEM structure using human TASK-1 protein identical to that used previously for crystallisation (Met1-Glu259).

We were able to resolve this structure to 3.1 Å resolution (**Figure 1, Supplementary Table S1 and Supplementary Figure S1**) and the overall structure was found to be consistent with previous biochemical and crystallographic structures of TASK-1. Superimposition of the cryoEM and crystal structures revealed very similar conformations (**Figure 1a-c**). The channel forms a domain-swapped homodimer with four transmembrane helices (M1-M4) in each subunit, a pseudotetrameric selectivity filter similar to other K2Ps, and an extracellular Cap domain that lacks the intermolecular disulphide bridge between chains found in most other K2Ps^29^. Even though the distal C-terminus beyond A251 is not resolved as well as in the crystal structure, the architecture of the X-gate (^243^VLRFMT^248^) is clearly visible and conserved, thus confirming the structural integrity of this gating motif. Cholesterol hemisuccinate was also identified bound to the same external clefts as seen in the TASK-1 crystal structure, i.e., next to the X-gate between the M1, M4 and M2 helices with a second site in a higher cavity along the M4 helices.

**Figure 1:**
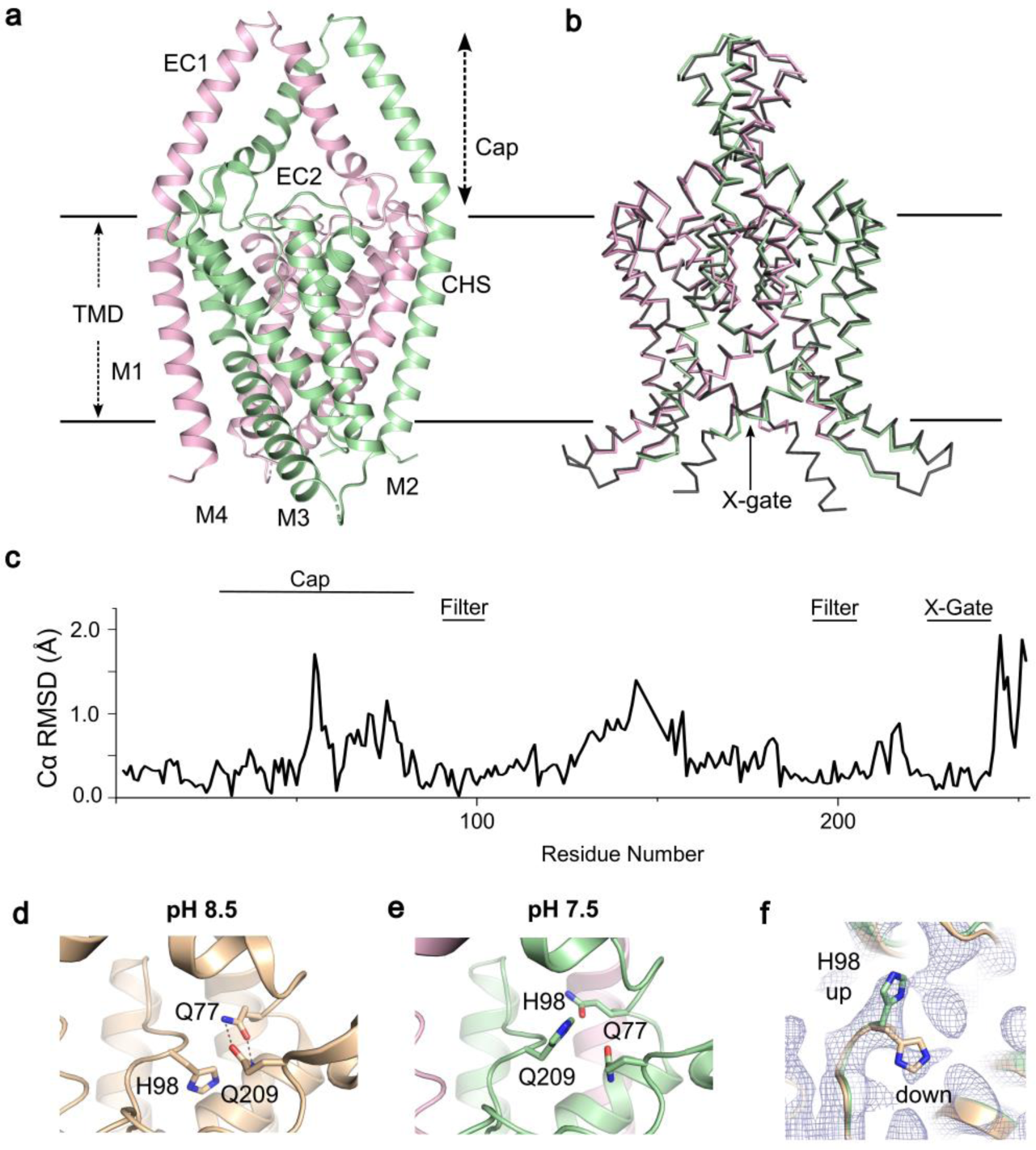
CryoEM structure of TASK-1. a. Cartoon showing the Cryo-EM structure of TASK-1, with A chain in green and B chain in pink. **b.** Superimposition of TASK-1 and TASK-1 crystal structure 6RV2 **c.** RMSD plot of the TASK1 cryoEM structure compared to the crystal structure (6RV2). **d.** H98 conformation of TASK-1 at pH 8.5 from 6RV2. **e.** H98 conformation at pH 8.5. **f.** Dual rotamer density of H98 with the upwards conformation shown and density for the down conformation clearly visible.

### TASK-1 pH-Sensor

Both TASK-1 and TASK-3 channels are sensitive to inhibition by extracellular H^+^ within the physiological range. The crystal structure of TASK-1 was originally solved at pH 8.5 with the pH-sensor, H98 presumably in the activated state where it points down to stabilise the region behind the filter via H-bond formation involving a water molecule and other side-chains within this region^17^ (**Figure 1d**). Similar networks of interactions have also been observed in other K^+^ channels stabilising their K^+^ selective filters in active, conductive conformations^26^. The H98 side chain is also adjacent to two glutamines (Q77 and Q209) at pH 8.5 in the crystal structure conformation with Q77 H-bonding with Q209, and mutations in this network of residues have been shown to alter TASK-1 pH sensitivity^17^.

Interestingly, despite the high degree of structural alignment in most other regions of the structure, the conformation of this pH-sensor in the new cryoEM structure, determined at pH 7.5, is markedly different to that previously found at pH 8.5 (**Figure 1e**). At pH 7.5, H98 adopts an alternative rotamer conformation and there is a disruption of the hydrogen bonds between Q77 and Q209. However, some density is still also observed for the downward rotamer (**Figure 1f**). This is consistent with the pKa for TASK-1 pH sensitivity and indicates that at pH 7.5 both conformations may exist. This therefore suggests that the first stages of pH sensitivity involve protonation of H98 which induces an upwards movement and disruption of the network of interactions between H98- Q77-Q209.

### CryoEM Structure of TASK-3

To investigate the structure of the related human TASK-3 channel we used an almost identical expression construct (Met1-Glu259) combined with a similar structural approach to determine a structure to 3.3 Å resolution (**Figure 2, Supplementary Table S1 and Supplementary Figure S2**). In TASK-3, the X-gate sequence motif (^243^VLRFLT^248^) is slightly different to that of TASK1 (^243^VLRFMT^248^). The overall fold of this TASK-3 structure is very similar to that of TASK-1 and almost identical to other structures of TASK-3 recently resolved by cryoEM^21^.

**Figure 2:**
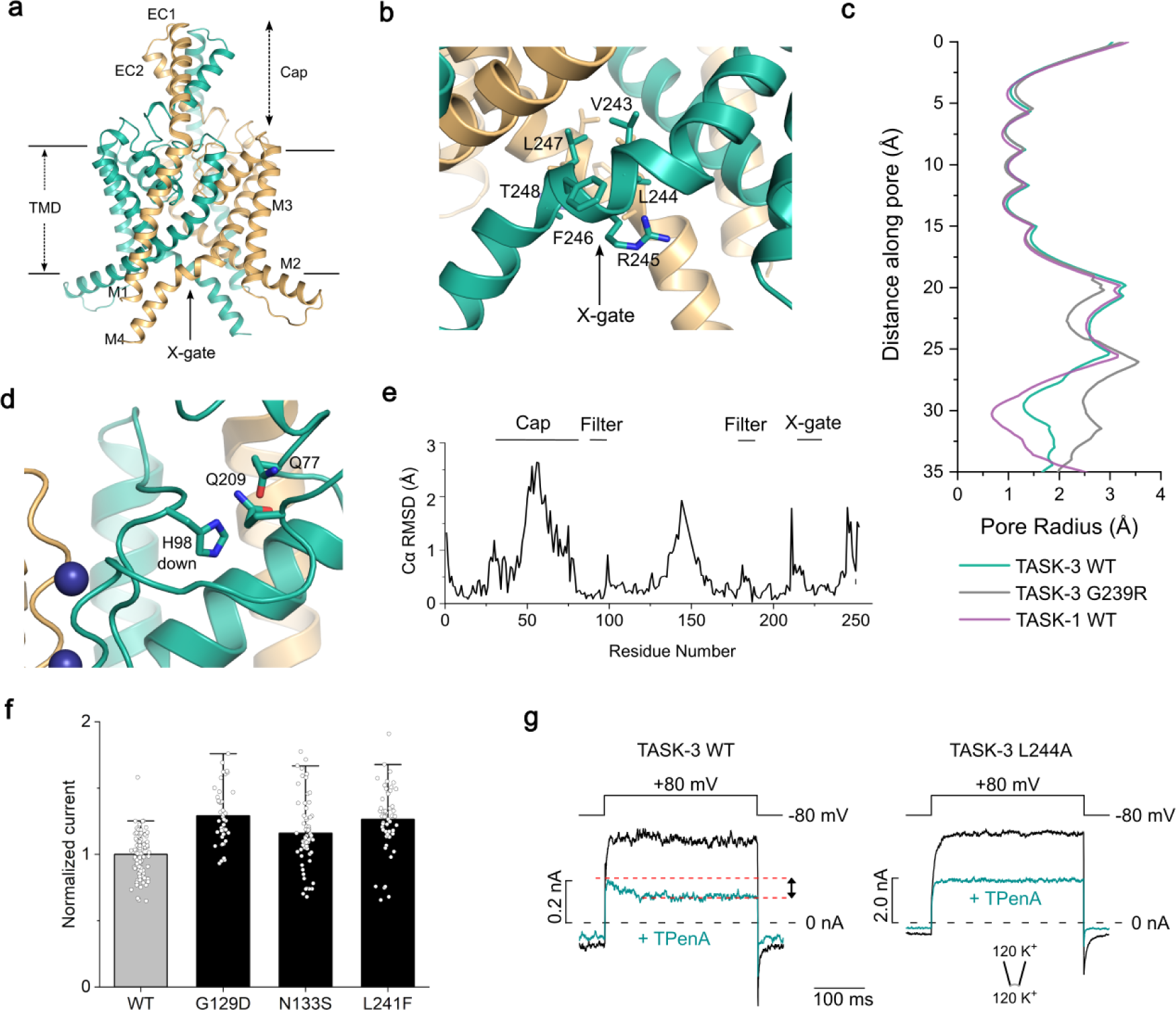
CryoEM structure of TASK-3. a. Cartoon showing the structure of TASK-3, with chain A in gold and chain B in teal. **b.** The X-gate environment of TASK-3 **c.** Pore radius profile of WT TASK-3 and the other structures determined in this study. **d.** H98 downward conformation. **e.** RMSD plot of TASK-3 versus TASK-1 **f.** Whole-cell recordings of WT TASK-3 (*n=95*) and TASK-3 with ‘DDSA’ mutations: G129D (*n=42*), N133S (*n=60*), and L241F (*n=51*). **g**. Current-voltage responses measured in inside-out patches from *Xenopus* oocytes expressing WT or L244A mutant TASK-3 channels in symmetrical high K^+^ at pH 7.4 in the absence (black) and in the presence of 1 µM TPenA (teal). The dotted red line indicates the extent of block by TPenA seen in WT TASK1 when the X-gate opens upon voltage-activation. This is not seen in the more active L244A mutant channel.

Together, these structures therefore confirm that TASK-3 possesses a structurally conserved lower X-gate motif where the distal end of the M4 helices kink and block the entrance to the inner cavity (**Figure 2a-e**). Similar to TASK-1, cholesteryl hemisuccinate was also found bound to the same regions next to the X-gate between the M1, M2 and M4 helices, with a second site in a higher cavity along the M4 helices. The pH-sensor (H98) is also found within the first filter motif (TIGYGH) in TASK-3, but in this structure solved at pH 7.5, only the downward facing rotamer was observed (**Figure 2d**). However, this is consistent with the slightly reduced H^+^ sensitivity of TASK-3 which is considerably more active at pH 7.5 than TASK-1 and so would be expected to be in the stable, conductive conformation.

The orientation of H98 is also consistent with structures of TASK-3 recently resolved by Lin *et al.* at pH 7.4^21^. However, in that study, the authors also solved a structure of TASK-3 at pH 6.0 which revealed much larger rearrangements of the filter backbone in this region. Intriguingly, these larger changes at pH6.0 could only be observed at low concentrations of K^+^, yet they share some similarity with a structure of TWIK-1 solved at pH 5.5 which suggest that protonation of a histidine at this position adjacent to the filter might induce a much larger rearrangement of the backbone to constrict the outermost part of the filter itself^20^.

### Conserved functional role of the X-gate in TASK-3

As found in TASK-1, the residues that surround the X-gate in TASK-3 are also involved in inter- and/or intra-subunit interactions known to influence channel gating in both channels^30^. For example, in a recent study, a number of disease-causing mutations in TASK-1 were shown to destabilize the X-gate, increase channel open probability (*Po*) and reduce GPCR-sensitivity^11^. To examine whether this X-gating mechanism is functionally conserved in TASK-3, we therefore examined the activity of several equivalent mutations in TASK-3. We found that, similar to their activatory effect in TASK1, the GoF DDSA mutations (G129D, N133S, and L241F) which destabilise the X-gate in TASK1 also increased whole-cell currents for TASK-3 (**Figure 2f**). The magnitude of this increase was less than that found in TASK-1, but this is probably due to the intrinsically higher open probability of WT TASK-3 to begin with. Indeed, it has previously been shown that a similar mutation of N133 in TASK-3 directly increases single channel open probability^31^ therefore supporting our observations that these mutations have an activatory effect due to the overall conservation of the X-gating motif in TASK-3.

Interestingly, we also found that functional measurements of macroscopic WT TASK-3 currents in excised patches using the universal K^+^ channel blocker tetrapentylammonium (TPenA) showed a slow component of inhibition (about 25 % of the total block) upon voltage activation. This block resembles the time course of inhibition seen with similar slow open channel blockers in classical voltage-gated K^+^ channels^32^. Furthermore, for TASK-3 channels, it was recently shown that voltage activation is achieved through a voltage-dependent SF gate^28^. These results therefore suggest that voltage activation also opens the lower X-gate in TASK-3, allowing TPenA to reach its binding site in the inner cavity and is comparable to the mechanism recently described for the related TALK-2 K2P channel^28^. To further examine this mechanism for TASK-3, we used a mutation at the X-gate (L244A)^17^ that has previously been shown to open the X-gate in TASK-1 and therefore expected to reduce this effect. Consistent with this, we found that the L244A mutation abolished this time-dependent component of blocker inhibition (**Figure 2g**). This not only supports a clear role for the X-gate in the gating of TASK-3 channels, but also highlights a degree of allosteric coupling between the filter and X-gate in these channels.

### Structure of the G236R TASK-3 mutation associated with KCNK9 Imprinting Syndrome

Both gain and loss of function mutations in TASK-3 lead to the neurodevelopmental disorder, KIS^15^. The most frequent mutation associated with this disorder is G236R on the M4 helix that results in significantly reduced channel activity. The precise mechanism by which this mutation affects channel activity is unclear because the mutant channels are not completely inactive and can be partially reactivated by TASK-3 activators such as flufenamic acid and terbinafine^33, 34^. We therefore used a similar approach to determine the structure of the G236R mutant channel to 2.5 Å resolution (**Figure 3, Supplementary Table S1 and Supplementary Figure S3**). Overall, the structure of this mutant channel was very similar to WT TASK-3, but with the mutant arginine side-chain from each monomer directly facing into the inner cavity to line the pore (**Figure 3b**).

**Figure 3:**
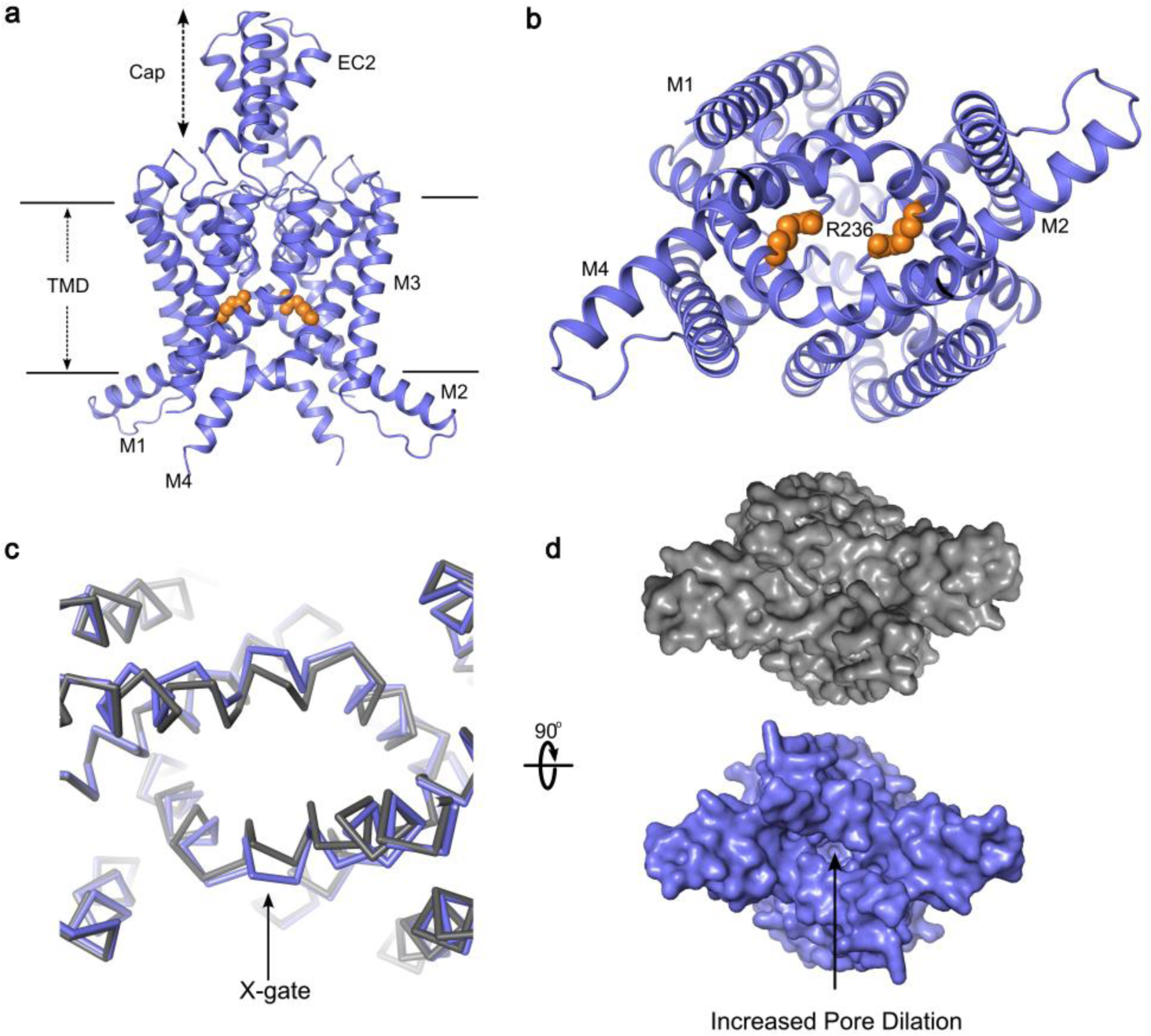
Cryo-EM structure of TASK-3 G236R. a, Cartoon showing the structure of TASK-3 G236R, with the two chains of the homodimer shown in blue. **b.** The position of R236 within the inner cavity is shown as orange VdW spheres. **c,d.** Overlay of TASK-3 (gray) and TASK-3 G236R (blue) highlighting the additional movement in the X-gate (expanded in panel d).

Intriguingly, it has also previously been shown that introduction of a negative charge into this position (G236D) produces an activatory effect^35^, and so it is likely that these positive charges directly interfere with the movement of K^+^ to reduce permeation through the channel thereby explaining the loss of function phenotype associated with this mutation. In addition, these two arginine residues are in relatively close proximity to each other within the inner cavity and we observed small changes in the local environment of the X-gate (**Figure 3c-d**). This movement causes the M4 helices to move apart in a direction similar to that seen in simulations of pore-opening in TASK-1 mutant channels^11^, and so may provide insight into the mechanisms which allow the aperture within the X-gate to expand during channel opening.

### Summary and Conclusions

Overall, our results provide further insight into the structure of both TASK-1 and TASK-3 channels and demonstrate that the X-gating mechanism is a highly conserved structural gating motif within the TASK channel family that, like many other K^+^ channels, also exhibits a degree of allosteric coupling between the lower gate and the filter gate. The alternate conformations of the H98 residue we observe in TASK-1 also provide evidence for the first changes that are likely to occur after protonation of this residue and suggest how flipping of the H98 side-chain may impact gating within the filter itself.

A structure of TWIK-1 at pH 5.5 reveals much larger rearrangements of the backbone that constrict the outermost part of the filter itself (**Figure 4**). Also, a very recent cryoEM structure of TASK-3 was solved at pH 6.0 with low K^+^ (5mM) which included much larger rearrangements of the filter backbone in this region^21^. In that structure of TASK-3, H98 has reoriented (**Figure 4c**), but is not in a fully upward conformation compared to H98 in the TASK-1 structure we present here at pH 7.5 (**Figure 4a**). Other key residues within the filter of TASK-3 also undergo large rearrangements at pH 6.0 and low K^+^, changes that are not seen in high K^+^. These include reorientation of Y96 and F202 in the P1 and P2 filter motifs from their positions behind the filter to restrict the pore opening, whilst in TWIK-1 at pH 5.5, H122 (corresponding to H98 in TASK channels) moves dramatically upwards, restricting the pore (**Figure 4d**).

**Figure 4.**
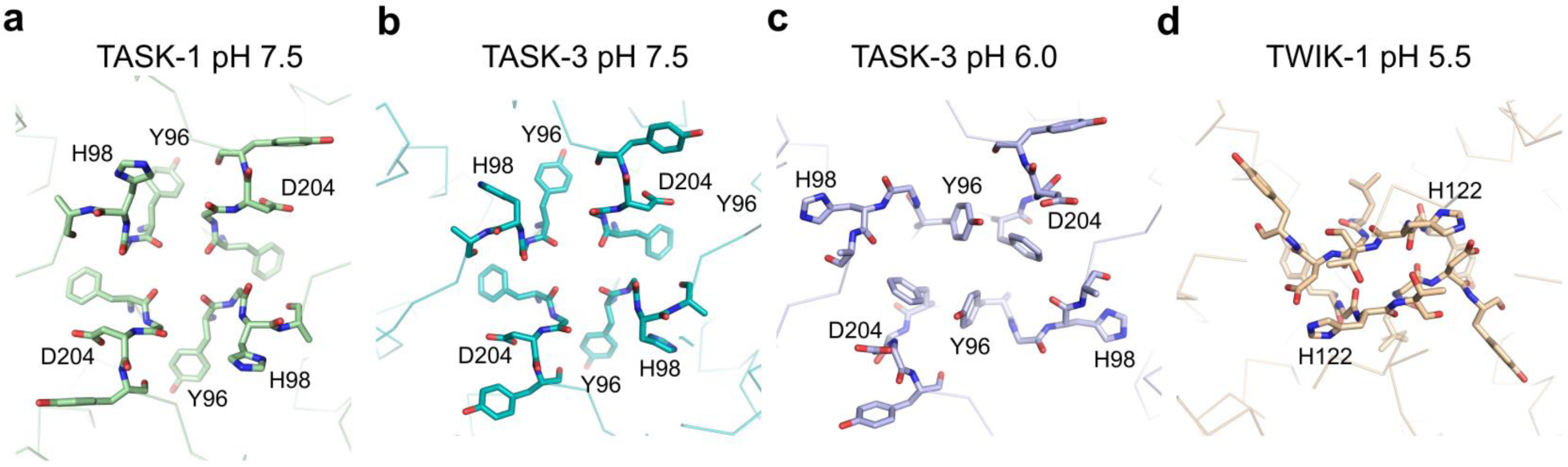
**Comparison of pH-dependent changes in the filters of TASK and TWIK channels. a**: TASK-1 at pH 7.5, H98 pointing upwards, D204 down (this study), **b**: TASK-3 at pH 7.5, H98 and D204 down, (this study) **c**: TASK-3 pH 6.0 low K^+ 21^, H98 reoriented downward D204 up, with Y96 and F202 pointing upwards, blocking the pore, **d**: TWIK-1 at pH 5.5 with major rearrangements in the loops, restricting the pore, H122 pointing up, D230 up.

However, it has been shown in other K2P channels, and in other K^+^ channels, that only small structural movements in the filter are required to affect the conformational dynamics of this region and influence K^+^ permeation^24–27^. Whether the small movement we observe in TASK-1 is sufficient to affect permeation by itself remains to be determined and instead may just represent the first stage of larger movements that occurs at more extreme pH values. It will therefore be interesting to examine whether such larger changes, especially in low K^+^, are physiologically relevant and necessary to gate the channel, and how these different conformational changes and their relative dynamics are associated with changes in the rate of K^+^ permeation through the filter itself.

Either way, it is clear that these new structures of TASK-1 and TASK-3 now expand the conformational landscape available for TASK channels as druggable targets and increase our insight into the mechanism of pH-sensing as well as a potential mechanism for coupling between the filter and the X-gate in TASK channels. The positively charged lining of the pore in the disease-causing G236R TASK-3 mutant also provides a clear structural explanation for the loss of function associated with this recurrent mutation and combined with more structural information for WT TASK-3 also now provide a robust structural framework for the development of activators against TASK-3 and other disease-associated mutations.

## Acknowledgments

This work was supported by grants from the Biotechnology and Biological Sciences Research Council and Medical Research Council to S.J.T, S.N and KEJR (BB/T002018/1, BB/S008608/1 and MR/W017741/1) and from the Deutsche Forschungsgemeinschaft to M.S. (SCHE 2112/1-2) and T.B (BA 1793/6-2) as part of the Research Unit FOR2518, DynIon. We would also like to acknowledge contributions from David Speedman and other members of the Centre for Medicines Discovery, Oxford during the early stages of this project and to Kathryn Smith for her assistance.

## Author contributions

S.J.T., S.N., P-R.H and K.E.J.R. conceived/designed the principal elements of the study. All authors generated, analyzed or interpreted data or generated materials. P-R.H, S.J.T, and K.E.J.R drafted the manuscript and all authors contributed to the final version.

## Conflicts of interest

The authors declare no competing financial interest.

## Data Availability

All data within this study are included in the article and/or the Supplementary Information, and materials are available upon request. The cryoEM model and maps for these structures have been deposited in the Protein Data Bank and the EMDB database, respectively, under the following accession codes:

**TASK-3**: 9G9V [https://doi.org/10.2210/pdb9G9V/pdb], and EMD-51158 [https://www.ebi.ac.uk/pdbe/entry/emdb/EMD-EMD-51158].

TASK-3 (G239R): 9G9W [https://doi.org/10.2210/pdb9G9W/pdb],

EMD-51159 [https://www.ebi.ac.uk/pdbe/entry/emdb/EMD-51159]. **TASK-1**: 9G9X [https://doi.org/10.2210/pdb9G9X/pdb] and EMD-51160 [https://www.ebi.ac.uk/pdbe/entry/emdb/EMD-51160].

## Materials and Methods

### Cloning and expression of TASK-3^Met1–Glu259^ for structural studies

The human *KCNK9* gene (GenBank ID 51305), encoding the TASK-3 protein residues M1 to E259, was subcloned into a modified pFastBac vector containing a C-terminal HRV 3C protease site, decahistidine and FLAG tags^17^. This construct was used for WT TASK-3 expression. TASK-3^G236R^, was subcloned into the pFB-CT10HF-LIC vector and comprised residues M1 to A265, this also contained a TEV protease site followed by a decahistidine tag and a FLAG tag. Baculoviruses were generated by transformation into DH10Bac cells and transfected into *Spodoptera frugiperda* (Sf9) cells (Thermo Fisher Scientific, 11496015) and amplified twice. Large scale expression was done by infecting Sf9 cells at a density of 2 × 10^6^ cells per mL and a 5 % virus ratio and grown at 27°C for 72 h. Cell were harvested by centrifugation at 900 g for 10 min and frozen in liquid nitrogen and stored at −80 °C prior to purification.

### Purification of TASK-1 and TASK-3

TASK-1 and TASK-3^G236R^ proteins were purified as previously described for TASK-1^17^, with the exception that TASK-1 was purified in *n*-undecyl-β-d- maltopyranoside (UDM) (Anatrace) instead of *n*-decyl-β-d-maltopyranoside (DM). The UDM concentrations throughout purification were 1% w/v for solubilisation, 0.09% for wash, elution and desalting buffers, and 0.06% for size exclusion buffer. The TASK-3 protein was purified by breaking the cells in 40 ml breaking buffer (50 mM HEPES pH 7.5, 200 mM KCl, 5% v/v glycerol) using an EmulsiFlex- C5 high-pressure homogenizer (Avestin). For solubilisation, 1% DM and 0.1% CHS Tris salt was added and rotated for 1 h at 4°C. The solubilised fraction was separated from cell debris by centrifugation at 35,000 *g* for 1 h at 4 °C and the supernatant then collected. To collect the histidine tagged protein, 0.5 ml Talon resin (Clontech) per initial litre of cell culture was added, as well as imidazole at pH 8.0 to a final concentration of 5 mM. Following incubation for 1 h at 4 °C, the Talon resin was collected and washed with 15 column volumes of wash buffer (50 mM HEPES pH 7.5, 200 mM KCl, 5% v/v glycerol, 20 mM imidazole pH 8.0, 0.24% w/v DM, 0.024% w/v CHS). The resin was transferred to a 15 ml centrifuge tube and the volume adjusted to 2 column volumes using wash buffer. Hexahistidine tagged HRV 3C protease and hexahistidine tagged PNGaseF were added to cleave the protein off the column, 150 µg and 50 µg per ml resin, respectively, and incubated for 16-20 h at 4 °C. The flow through was collected and concentrated to 500 µl and injected onto a Superose 6 Increase 100/300 GL column (Cytiva) in gel filtration buffer (20 mM HEPES pH 7.5, 200 mM KCl, 0.12% w/v DM, 0.012% w/v CHS). Fractions containing TASK-3 were pooled and concentrated with a 50 kDa molecular weight cut off spin concentrator (Cytiva).

### Cryo-EM grid preparation and data collection

For TASK-1 (3.6 mg mL^-1^) and TASK-3 (8 mg mL^-1^), 3 μL were adsorbed to glow-discharged holey carbon-coated grids (Quantifoil 300 mesh, Cu R1.2/1.3). were then blotted for 3 to 6 s at 100% humidity (4 °C) and frozen in liquid ethane using a Vitrobot Mark IV (Thermo Fisher Scientific). Data were collected in counted superresolution bin 2 mode on a Titan Krios G3 (FEI) operating at 300 kV with a BioQuantum imaging filter (Gatan), and K3 direct detection camera (Gatan) at ×105,000 magnification, physical pixel size of 0.832 Å. Collection of TASK-1 data was conducted at a total dose rate of 41.173 e^−^/Å^2^ and 17,288 movies were collected in total. The grids used for collecting this TASK-1 data were prepared after adding 1.3 µm ONO-RS-082, a TASK-1 activator, but density corresponding to this small molecule activator could not be identified in the Coulomb potential map possibly due to the relatively low solubility of the compound and low final concentration that was used. The TASK-3 data were collected with a total dose of 39.56 e^−^/Å^2^ and 21,364 movies were collected in total. For TASK- 3^G236R^ (6 mg mL^-1^) the same process was conducted using holey gold grids (Au-Flat 300 mesh, R1.2/1.3, 45 nm gold film) with a total dose rate of 40.877 e^−^/Å^2^ and 20,627 movies.

### Image processing

Initial micrograph processing of all three datasets was conducted using the SIMPLE pipeline^36^, using SIMPLE preprocess_stream for motion correction and patched contrast transfer function (CTF) estimation. Particles were picked from a subset of each dataset, and extracted in SIMPLE. All further processing was done in cryoSPARC^37^. Motion corrected micrographs were imported to cryoSPARC and CTF estimation was done. The initial particles extracted in SIMPLE were imported and subjected to 2D classification rounds and ab-initio reconstitution to yield a volume which templates for picking were generated. Particle repicking from the full dataset was done using the template picker within cryoSPARC. All further processing was done in cryoSPARC unless stated otherwise. For WT TASK-1, 8,116,956 particles were extracted and underwent one round of unmasked 2D classification, followed by four rounds of masked 2D classification and junk particles were removed. The resulting set of 1,076,556 particles were classified by five C1 ab initio classes and after heterogenous refinement, three of these were selected for further processing. After one round of masked 2D classification, the remaining 680,017 particles were classified in three C2 ab initio classes. After heterogenous refinement and homogenous refinement of one class, the particles (358,967) were exported to RELION-5^38^, using the csparc2star.py script within UCSF pyem and subjected to Bayesian polishing. The polished particles were run through cryosieve^39^ and particle stacks were imported back into cryoSPARC^37^. To find the highest resolution stack, each stack was subjected to one round of 2D classification, ab initio reconstruction in C1 followed by homogenous and heterogenous refinements, both in C2. The final particle set (117,627) yielded a map of 3.13 Å resolution, based on the FSC = 0.142 criteria. For TASK-3, 12,215,148 particles were initially extracted, which underwent two rounds of 2D classification. Subsequently, five C1 ab initio classes were generated from 1,904,926 particles. Following heterogeneous and non uniform refinement, 597,895 particles were selected for further analysis. After 2D classification, ab initio reconstruction in C2 with five classes, and heterogenous refinement applying C2 symmetry, one class with 286,131 particles was subjected to homogenous and non uniform refinements in C2. These particles were then exported to RELION and Bayesian polished, run through cryosieve and particle stacks were imported back into cryoSPARC, where they were processed as described above for TASK-1. The final C2 map from 93,760 particles was resolved to 3.32 Å. For TASK-3^G236R^, 8,932,733 particles were extracted and subjected to one round of unmasked 2D classification, followed by five rounds of masked 2D classification, resulting in a set of 2,014,115 particles. These were sorted into five C1 ab initio classes and underwent heterogenous refinement. Three classes, containing junk and lower resolution particles, were discarded and the remaining two were classified in 2D to remove further junk particles, resulting in a set of 1,145,364 particles. They were then classified into three C2 ab initio classes and after heterogenous refinement, one class underwent heterogenous refinement. These particles were Bayesian polished in RELION and narrowed down using cryosieve. Particle stacks were imported to cryoSPARC and went through 2D classification, ab initio reconstruction of one class in C1, and refinements in C2, as described for the TASK-1 map. The final particle set of 124,588 particles yielded a map at 2.48 Å.

### Model building, fitting and refinement

Both TASK-3 models were manually built in the map on Coot to generate the initial model. After which the models were manually readjusted using COOT^40^ and refined using *phenix.real_space_refine*^41^. For TASK-1, the X-ray crystal structure (PDB ID: 6RV2) was used as a starting model and manually fitted into the map in COOT. Final models were run through ISOLDE^42^ and refined in phenix.real_space_refine using the generated ISOLDE models as references.

### Electrophysiology

The WT TASK-3 gene (*KCNK9*) and human TRAAK (*KCNK4*) genes were subcloned into a plasmid vector (pFW) *Xenopus laevis* oocyte transcription and expression. The mutations were introduced through site-directed mutagenesis and unless stated otherwise the two-electrode voltage clamp recordings were performed as previously described^11^. Vector DNA was linearized with NheI or MluI and cRNA synthesized *in vitro* using the SP6 or T7 AmpliCap Max High Yield Message Maker Kit (Cellscript, USA) or HiScribe® T7 ARCA mRNA Kit (New England Biolabs) and stored at -20°C. TEVC recordings were performed in *Xenopus laevis* oocytes at room temperature. Oocytes were injected with 1 ng of cRNA for WT or mutant TASK-3 and incubated for 2 days at 18°C. Microelectrodes were fabricated from glass pipettes, back-filled with 3 M KCl, and had a resistance of 0.2 - 1.0 MΩ. For inside-out patch-clamp recordings oocytes were incubated in a solution containing (mM): 54 NaCl, 30 KCl, 2.4 NaHCO3, 0.82 MgSO4 x7H2O, 0.41 CaCl2, 0.33 Ca(NO3)2 x4H2O and 7.5 TRIS (pH 7.4 adjusted with NaOH/HCl) for 1-7 days before use. Inside-out patch-clamp recordings were performed at room temperature with patch pipettes were made from thick-walled borosilicate glass with resistances of 0.3 - 0.5 MΩ (tip diameter of 15-25 µm) and filled with a pipette solution (in mM): 120 KCl, 10 HEPES and 3.6 CaCl2 (pH 7.4 adjusted with KOH/HCl). Intracellular bath solutions and compounds were applied to the cytoplasmic side of excised patches via a gravity flow multi-barrel pipette system. Intracellular solution had the following composition (in mM): 120 KCl, 10 HEPES, 2 EGTA and 1 Pyrophosphate (pH adjusted with KOH/HCl). Currents were recorded and sampled at 10 kHz or higher and filtered with 3 kHz (-3 dB) or higher as appropriate.

## Supplementary Information

**Table 1.** Cryo-EM data collection, refinement and validation statistics.

|  | **TASK-3** | **TASK-3 (G236R)** | **TASK-1** |
| --- | --- | --- | --- |
|  | **9G9V EMD-51158** | **9G9W EMD-51159** | **9G9X EMD-51160** |
| **Data collection and processing** |  |  |  |
| Magnification | 105,000 | 105,000 | 105,000 |
| Voltage (kV) | 300 | 300 | 300 |
| Electron exposure (e-/Å<sup>2</sup>) | 39.56 | 40.877 | 41.173 |
| Defocus range (µm) |  |  |  |
| Pixel size (Å) | 0.832 | 0.832 | 0.832 |
| Symmetry imposed | C2 | C2 | C2 |
| Initial particles images (no.) | 12,215,148 | 8,941,912 | 8,116,956 |
| Final particle images (no.) | 93,760 | 124,588 | 117,627 |
| Map resolution (Å) | 3.32 | 2.48 | 3.13 |
| FSC threshold | 0.143 | 0.143 | 0.143 |
| **Refinement** |  |  |  |
| Initial model used | Manually | Manually | Crystal structure (6RV3) |
| Non hydrogen atoms | 4073 | 4089 | 3955 |
| Protein residues | 522 | 516 | 488 |
| Average *B* factors (Å<sup>2</sup>) |  |  |  |
| Protein | 79.30 | 58.75 | 40.48 |
| Ligand | 83.58 | 67.09 | 53.64 |
| r.m.s.d. |  |  |  |
| Bond lengths (Å) | 0.008 | 0.008 | 0.008 |
| Bond angles (°) | 1.140 | 1.111 | 1.143 |
| **Validation** |  |  |  |
| MolProbity score | 0.93 | 0.82 | 0.80 |
| Clashscore | 1.73 | 1.11 | 1.26 |
| Poor rotamers (%) | 0.53 | 0.00 | 0.00 |
| **Ramachandran plot** |  |  |  |
| Favoured | 98.46 | 99.61 | 99.17 |
| Allows (%) | 1.54 | 0.39 | 0.83 |
| Outlier (%) | 0.00 | 0.00 | 0.00 |

**Supplementary Figure S1.**
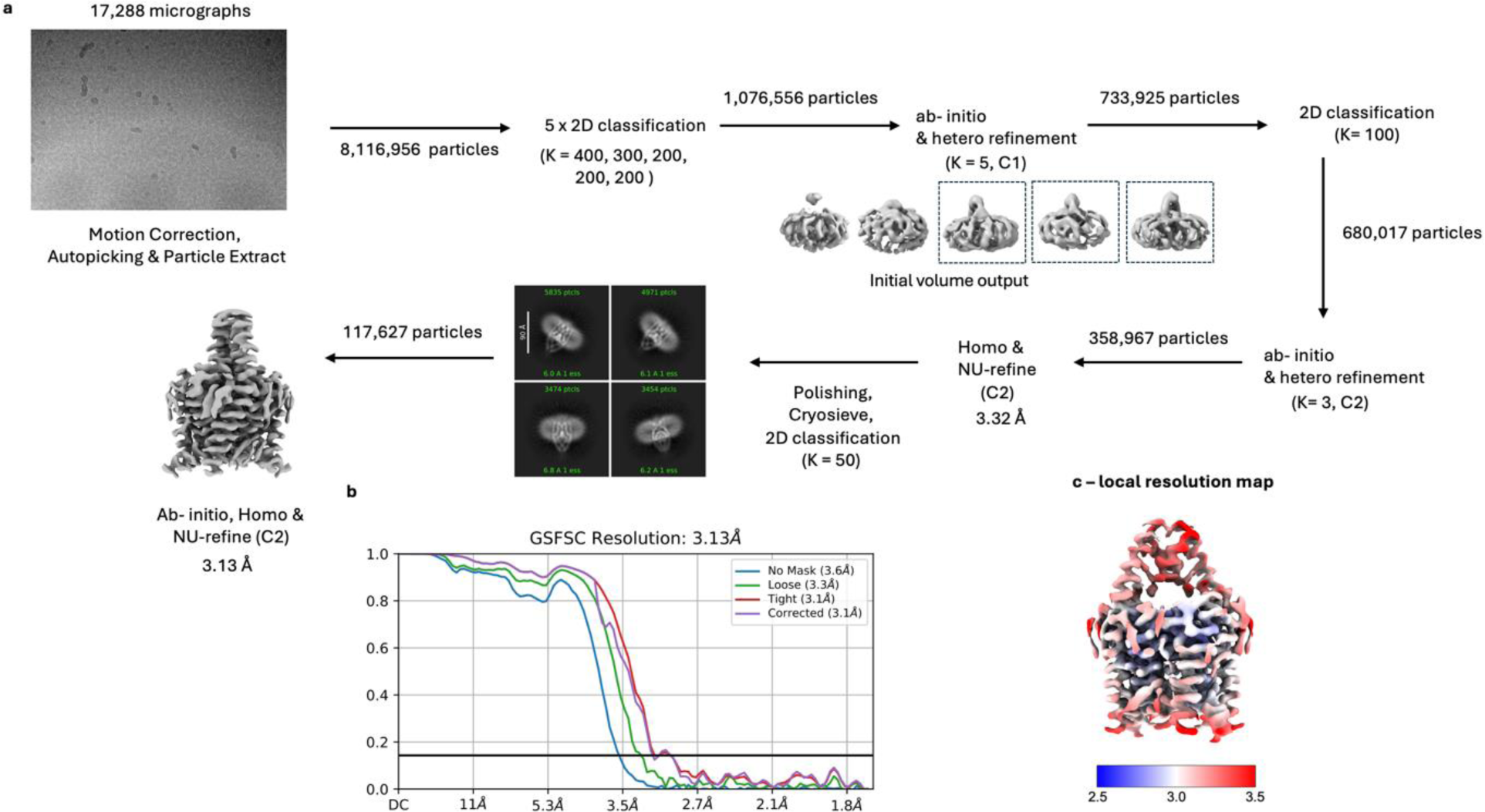
**Cryo-EM processing workflow for TASK-1 showing the local resolution map**. **a**, Image processing workflow for TASK-1. **b**, Gold-standard FSC curve used for global-resolution estimates within cryoSPARC. **c**, Local-resolution of reconstructed map as determined within cryoSPARC.

**Supplementary Figure S2.**
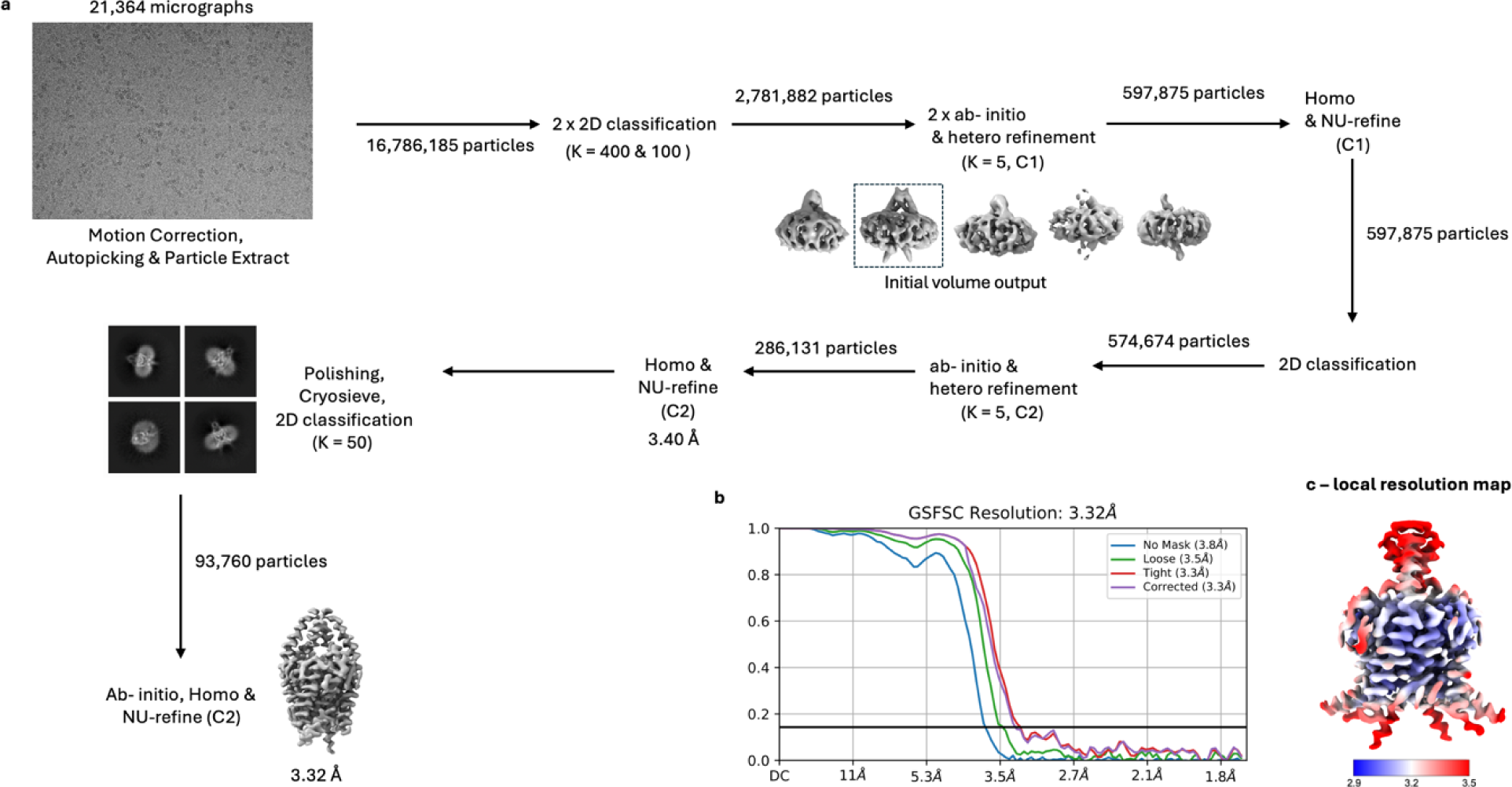
**Cryo-EM processing workflow for TASK-3 showing the local resolution map**. **a**, Image processing workflow for TASK-3. **b**, Gold-standard FSC curve used for global-resolution estimates within cryoSPARC. **c**, Local-resolution of reconstructed map as determined within cryoSPARC.

**Supplementary Figure S3.**
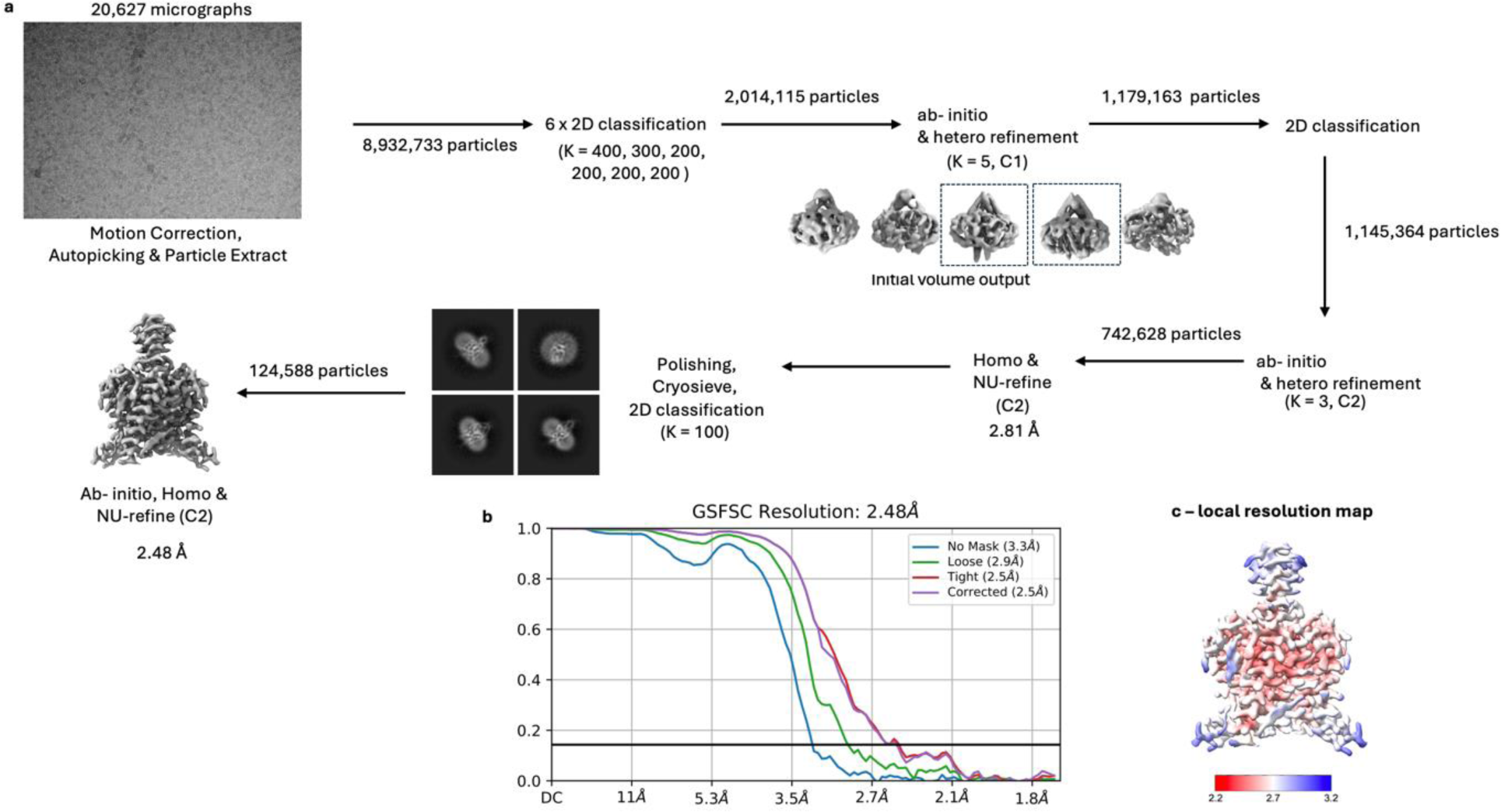
**Cryo-EM processing workflow for G236R TASK-3 showing the local resolution map**. **a**, Image processing workflow for G236R TASK-3. **b**, Gold-standard FSC curve used for global-resolution estimates within cryoSPARC. **c**, Local-resolution of reconstructed map as determined within cryoSPARC.

## Notes

### Competing Interest Statement

The authors have declared no competing interest.

